# Iterative Machine Learning for Classification and Discovery of Single-molecule Unfolding Trajectories from Force Spectroscopy Data

**DOI:** 10.1101/2023.08.08.552253

**Authors:** Vanni Doffini, Haipei Liu, Zhaowei Liu, Michael A. Nash

## Abstract

We report the application of machine learning techniques to accelerate classification and analysis of protein unfolding trajectories from force spectroscopy data. Using kernel methods, logistic regression and triplet loss, we developed a workflow called Forced Unfolding and Supervised Iterative Online (FUSION) where a user classifies a small number of repeatable unfolding patterns encoded as image data, and a machine is tasked with identifying similar images to classify the remaining data. We tested the workflow using two case studies on a multi-domain XMod-Dockerin/Cohesin complex, validating the approach first using synthetic data generated with a Monte Carlo algorithm, and then deploying the method on experimental atomic force spectroscopy data. FUSION efficiently separated traces that passed quality filters from unusable ones, classified curves with high accuracy, and identified unfolding pathways undetected by the user. This study demonstrates the potential of machine learning to accelerate data analysis, and generate new insights in protein biophysics.

## Introduction

Single-molecule force spectroscopy (SMFS) is a well-established analytical technique capable of generating unique insights into proteins^1,2^ and revealing detailed information about protein structure,^3^ mechanics,^4–6^ folding^7–9^ and function.^10–12^ SMFS can be carried out using a variety of force-based devices including magnetic tweezers, optical tweezers, centrifuge force microscopy or atomic force microscopy (AFM).^13^ Due to the comparatively stiff spring constants of typical cantilevers, AFM is most suitable for studying molecular systems^14^ in a high force regime (50 – 2500 pN) and remains a powerful tool for measuring the mechanical strength of protein-protein interactions, ^14^ protein unfolding transitions,^15^ and force-activated catch bonds. Single-cell analysis with the AFM is a closely related method that provides insights into cell surface composition, avidity of cell adhesion receptors, and information on cell mechanics with important connections to cell biology, biomedicine and therapies.

Typically, to measure protein-protein interactions under force, a binding receptor is covalently linked to a cantilever tip and the ligand is attached to a surface. The receptor and ligand are brought into contact and the cantilever is retracted. This generates a force-extension curve containing features that report molecular unfolding transitions and final dissociation of the molecular complex. Typically, the vast majority of AFM force-extension curves are not usable due to non-specific adhesion events, lack of interactions, or other spurious signals.^16^ Thousands of data traces are typically obtained using automated measurement routines, and this massive data set is then filtered, sorted and classified during post-processing and analysis. During data processing, some unwanted traces can be excluded by filtering for basic threshold features, such as minimum force value or a minimal distance from the surface for specific events (e.g., bond rupture or domain unfolding). Furthermore, contour length transformation is a valuable data analysis technique that maps the force-extension behavior into force-contour length space.^16,17^ By including control elements into AFM protein constructs (i.e., fingerprint domains), cross-correlation analysis of contour length histograms can be used to align and filter the data traces and identify those that capture valid single proteinprotein interactions. Nonetheless, it is difficult to set clear criteria a priori for classification of good and bad curves, and to separate different unfolding pathways for the same system. Since AFM-SMFS data is generally scarce and costly to produce, researchers may prefer to tolerate false positives rather than excluding all true negatives, and filters are usually set loosely. It remains the case that even with the best contour length analysis, in order to perform the technique to a high standard, there is a need to manually check, validate and classify thousands of data traces. This process is time-consuming and prone to errors and biases.^18^ This approach also relies on the user correctly identifying the repeatable unfolding patterns, potentially leaving rare pathways unidentified and lost.

Machine Learning (ML) can potentially address this challenge. ML has been used extensively for image enhancement^19^ and screening of topographical, phase and amplitude AFM imaging data. ^20–24^ However, its use for processing SMFS data curves is much more limited. A recent study^25^ trained a 1D Convolutional Neural Network (CNN)^26^ using a triplet loss function^27^ to generate an embedding space where curves containing single, double and multiple or no rupture events were separated. The authors demonstrated 65-70% accuracy in classifying a subset of data traces that were reliably and consistently classified by experts. However, in that study the dataset was class balanced, which is unrealistic for unprocessed AFM-SMFS datasets that typically contain a high proportion of negative (i.e. non-interaction) curves. That prior study furthermore did not examine classification of unfolding pathways in complex multi-domain systems, or address identification of pathways that were not previously identified by the user. If we could use ML as an iterative tool to accelerate the classification of SMFS curves, while enabling the discovery of novel unfolding pathways, we would accelerate biophysics research and enable the identification of rare molecular events that are currently invisible to the user.

In this work, we present a technique called Forced Unfolding and Supervised Iterative ONline learning (FUSION learning), based on a fixed pre-trained convolutional neural network (DenseNet121^28^) and kernel classification/embedding. FUSION learning iteratively separates meaningful traces from unspecific interactions (Layer 1) and generates an embedding space where different unfolding pathways are separated and visualized (Layer 2). Moreover, we show how covariance analysis can be used to detect previously unidentified unfolding pathways (Layer 3). We tested and validated our approach on a multi-domain XMod-Dockerin/Cohesin receptor-ligand system using synthetic force traces simulated with a Monte Carlo algorithm, and then on AFM-SMFS experimental data. We used this framework to accelerate the classification of unfolding trajectories and discover novel reaction pathways unknown to the user.

## Results and discussion

We used simulated and experimental force-extension data (as described in the Supporting Information, Methodology section) generated from the forced pulling process of a mechanostable XModule-Dockerin/Cohesin (XMod-Doc/Coh) complex (Fig. 1A) derived from a multi-protein cellulolytic complex from R. champanellensis (Rc).^29,30^ This molecular complex has been systematically studied using AFM-based single-molecule force spectroscopy previously by our group.^31,32^ The complex exhibits a characteristic dual binding mode behavior where three different data patterns can be assigned to different unfolding/dissociation pathways (P1, P2 and P3).^31^ P1 results in a high force rupture lacking XMod unfolding. Along pathway P2, the Ig-like XMod that stabilizes the complex unfolds. P3 meanwhile constitutes a low force rupture event lacking XMod unfolding. This complexity makes it challenging to filter and classify curves into the various pathway classes and presents a good opportunity to validate our FUSION learning approach. We used raw experimental data and also generated simulated force-extension traces using a dual-binding mode multi-state kinetic model with relevant parameters extracted from the experimental dataset.

**Figure 1:**
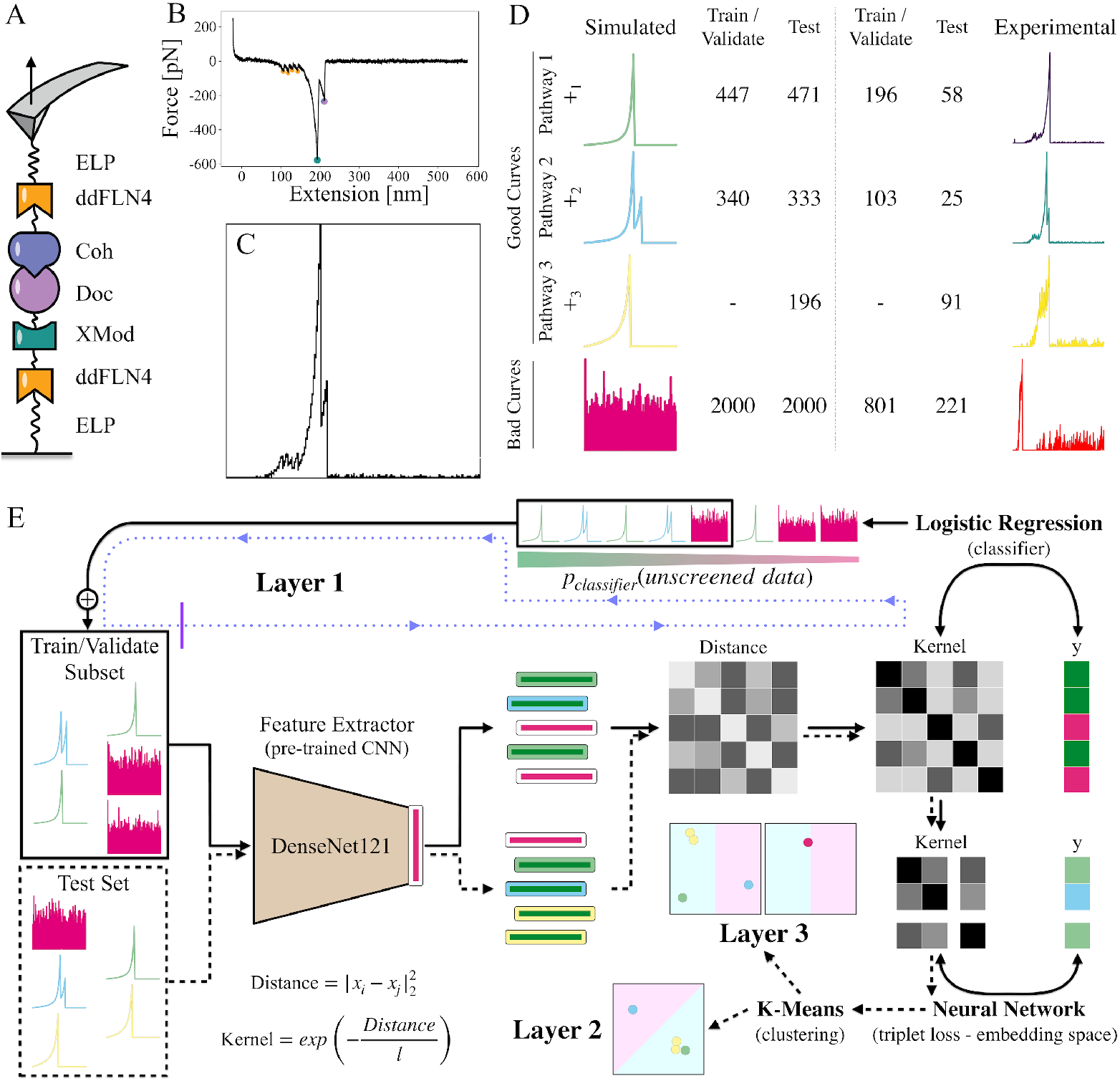
Summary of source data, preprocessing and FUSION learning algorithm. (A) Experimental configuration of multi-domain XMod-Dockerin/Cohesin complex. (B) Example of an experimental data trace. The colored dots highlight the rupture events of the individual domains color coded from panel A. (C) Data trace converted into an image through normalization to peak force value. (D) Summary of the datasets used in this work. Note that P3 curves are not included in the training and validation sets as a test case for identifying unknown pathways. (E) Overview of the FUSION learning workflow. The processed images are encoded with a pre-trained version of DenseNet121. The resulting features are used to calculate a Euclidean distance matrix, which is used to calculate a similarity matrix from a Gaussian kernel. This matrix is then used to fit: i) a binary classifier which sorts the unseen data, selects the best ones and lets the user check them before merging with the training and validation subset (layer 1); and ii) a triplet loss based neural network, which generates an embedding space where selected (layer 2) and unselected traces (layer 3) are clustered. The solid and dashed lines correspond to the screened (train/validate subset) and test data streams.

For ML training, the experimental dataset^31^ was divided into two sub-sets (Experimental, Fig. 1D). One subset was used for training and validating and the other for independent testing. For the simulated case, the Monte Carlo algorithm was used to generate two datasets, again one for training and validation and the other for testing, containing the same number of curves (Simulated, Fig. 1D). To simulate the presence of traces not containing any adhesion/interaction events, we generated 2,000 curves of normally distributed noise and added these curves to both simulated datasets. Additionally, in the experimental case all traces exhibiting the low force unbinding interaction (Pathway 3) were assigned to the test dataset, while in the simulated case such curves were simply removed from the train/validate set. This was done to test the capability of our workflow to detect an unselected curve class (i.e., a curve class unidentified by a user). We deliberately opted to exclude the curves labeled as Pathway 3 from the train/validate set to simulate two distinct scenarios. In the first one, we considered the possibility that Pathway 3 was intentionally left unlabeled by the user due to its perceived lack of significance in comparison to other labeled pathways. The second scenario aimed to simulate a situation where an unexpected or undetected change in the reaction conditions occurred, leading to the emergence of Pathway 3 at a later stage. This allowed us to explore the challenges associated with identifying and classifying an undetected pathway in such dynamic conditions. Furthermore, the decision to exclude these curves stemmed from their striking similarity to the high rupture forces observed in Dockerin/Cohesin interactions (i.e., Pathway 1) or to some unspecific interactions. This resemblance not only posed a challenge for ML-guided identification of Pathway 3 but also enhanced the realism of the task by requiring the model to discern subtle differences between similar features.

To pre-process the data, each force-extension curve (Fig. 1B) was down-sampled, inverted, trimmed and normalized by the maximum force (see Supporting Information). The obtained normalized traces were used to generate square 2D images (Fig. 1C), which were then encoded in vectors using the initial upstream convolution and pooling layers of DenseNet121,^28^ a pre-trained CNN (Fig. 1E). The DenseNet121 neural network was originally trained on the ImageNet dataset,^33^ which contains labelled examples of common objects. To use DenseNet121 as a feature extractor, we removed the last dense layer of the network, which is usually responsible for indicating the main object present in the input image. Even though DenseNet121 was never trained on AFM images or SMFS datasets, its ability to distill low-level attributes, such as contours and edges from image data are useful for our task. We deliberately chose to not fine-tune DenseNet121 during the training to: i) speed up the learning process, and therefore its iterability; ii) avoid overfitting, coming from the sparsity of data available; and iii) show the transferability of complex models that were pre-trained for a different application on AFM data.

The iterative ML algorithm (Fig. 1E) was initialized by showing an optimal artificial user a certain number of random sampled images of AFM traces (i.e., train and validate subset). These images were correctly labelled by the artificial user the same way they were labelled in the dataset (i.e., unspecific curve, or specific curve of Pathway class P1 or P2). Then, the corresponding feature vectors were used to generate a Gaussian kernel matrix (i.e., a similarity matrix), which was used as input to fit a logistic regression. After this first step, the binary classifier was run through the remainder of the unscreened curves in the dataset, ranking each curve according to its probability to be labeled ‘specific’ or ‘unspecific’ (i.e., no interactions or unusable curve), and feeding the best results to the user for another iteration. This gave to the user the possibility to accept or correct the predictions of the model, again on a restricted number of top results, increasing the Train/Validate subset dimension at each iteration.

At this stage (Layer 1), the model could only distinguish between specific and unspecific curves, but it could not yet separate different user-selected unfolding pathways (Layer 2). To enable pathway classification, an additional model was employed. First, the rows and columns corresponding to the unspecific curves were removed from the kernel matrix. Then, a dense neural network was trained on top of the similarity matrix by minimizing triplet loss. ^27^ Such loss functions are usually applied to train supervised models for image clustering. Here, we used it to generate an embedding space where the traces selected by the classifier could be visualized. If trained correctly, and after a sufficient number of iterations (i.e., enough training data), curves containing similar unfolding pathways should be located close to each other in the resulting embedding space. In addition, to measure the “goodness” of pathway separation in this space, we applied a k-means clustering algorithm on the embedded data. Note that, since the iterative screening by classification (Layer 1) was completely independent from this additional neural network, the user could decide at which iteration to generate the embedding space.

At this point, our triplet loss neural network (Layer 2) was not sufficient to successfully separate unfolding pathways that were not explicitly labeled by the user, due to overlapping of the different classes. For this reason, we next expanded our embedding space by adding a new dimension: the posterior covariance.^34^ This quantity, calculated from the Gaussian kernel matrix, can be seen as a measure of the distance between each test point to the data used during the training. Therefore, since data containing unlabeled classes were somewhat different from labeled ones, it would be possible to separate them and identify unknown unfolding pathways.

In the following two case studies, the Layer 1 (iterative classifier) was employed to screen a dataset (Fig. 1D, train/validate), while the Layer 2 (embedding) and Layer 3 (covariance) were validated using an independent test set containing an additional unfolding pathway (Pathway 3), which was not present in the other dataset (Fig. 1D).

To verify the applicability of our framework, we first employed it on synthetic data generated with a Monte Carlo algorithm^35,36^ (Simulated, Fig. 1D). This case study served as a proof-of-concept, where each good trace was clearly defined, separable and virtually noise free. The bad curves consisted of pure noise, simulating the absence of any molecular adhesion interactions.

The workflow was initialized by randomly sampling and labeling 4 curves from the train/validate dataset (Fig. 1D). Such traces constituted the train/validate subset (Fig. 1E) in the first iteration, which was used to train the logistic regression model for the first time. Then, the binary classifier was used to predict the probability of each trace that was not yet present in the subset (unscreened data, Fig. 1E) was a good curve (i.e. positive interaction). After that, the top-4 traces were selected and merged in the train/validate subset (Fig. 1E) for the next iteration. Note that, if some labels of those 4 curves were misclassified by the logistic regression, we corrected them, simulating a human supervisor who can distinguish between bad and good curves.

To evaluate the acceleration in the screening of a specific dataset (Layer 1), we report the percentage and the absolute number of good curves found at each iteration against the absolute and relative number of traces analyzed (Fig. 2A). We further plotted the theoretical performance of a naïve screening through the dataset (random sampling, Fig. 2A) as a baseline. The naïve approach corresponded to a naïve user analyzing the curves without the ability to reorder them according to the output probability of the binary classifier. For this reason, our baseline consisted in equalizing the percentage of good curves found with the percentage of curves analyzed. Similarly to a receiver operating characteristic (ROC) curve, the higher a curve-screening strategy was from the diagonal (i.e., the random sampling baseline), the higher the benefit of using FUSION learning.

**Figure 2:**
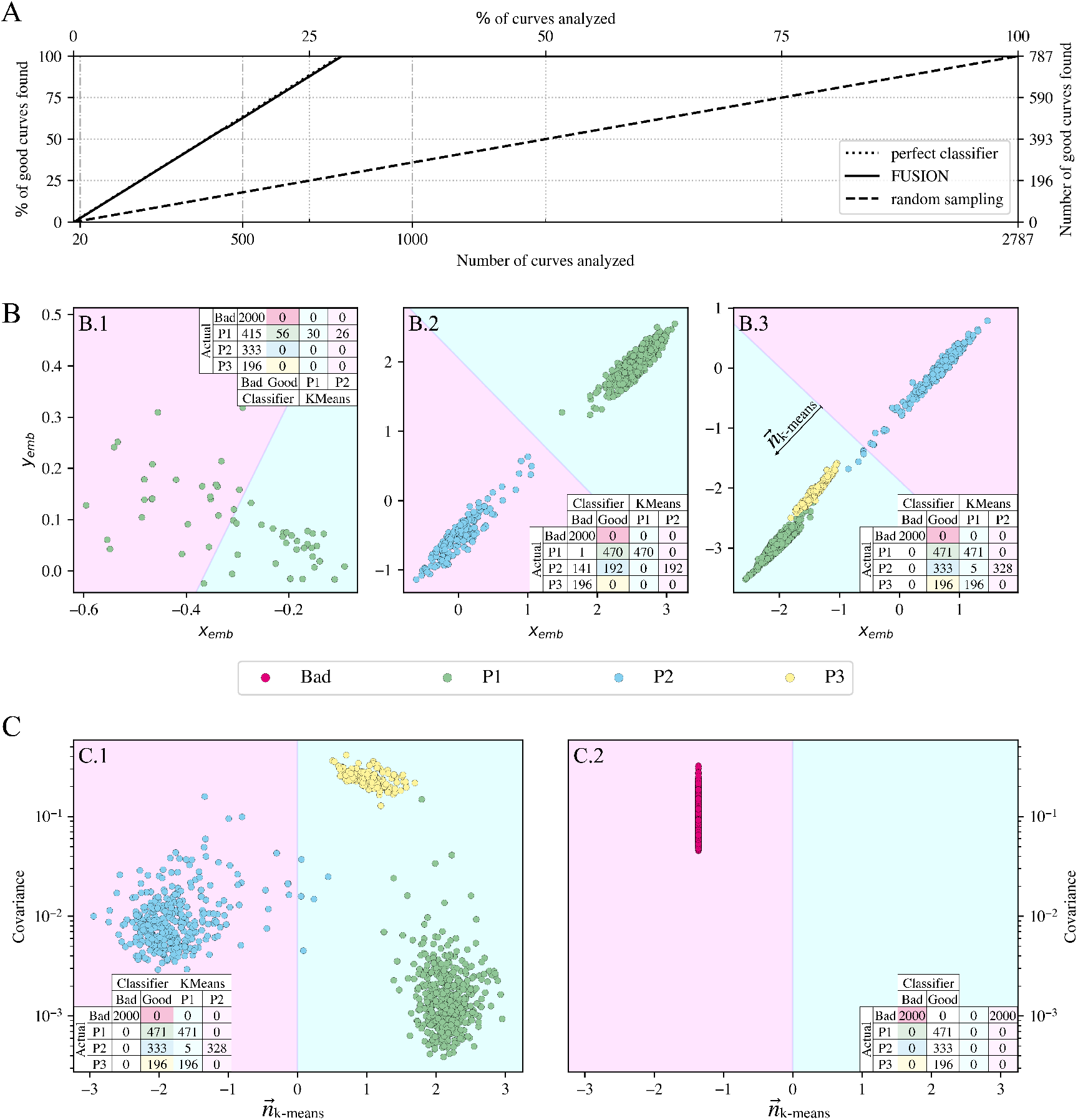
Results of the case study using simulated data. (A) Layer 1: iterative classification of good curves. The solid black line shows the result of our FUSION learning method. The dashed black line shows the theoretical random sampling. (B) Layer 2: embedding space generated at different iterations, after analyzing 20 (B.1), 500 (B.2) or 1000 (B.3) traces. The data shown come from an independent test set. Only the spectra classified as good are shown. The colors of the dots recall the true labels. The background coloring indicates the separation boundary of a k-means algorithm. (C) Layer 3: separation of selected (P1 & P2), unselected (P3) and bad curves by covariance and the output of the binary classifier (layer 1) at 1000 analyzed spectra. (C.1) contains the curves classified as good, while (C.2) contains the traces classified as bad.

Using our approach, all good curves (447 P1 curves + 340 P2 curves = 787 good curves) enclosed in the train/validate Monte Carlo dataset (n=2,787 curves total) were correctly sorted at the 199^*t*^*h* iteration (796 curves screened of which 787 were good, 9 bad). This means that we were able to identify all the useful curves after screening only 28.56% of the dataset. This result corresponded to an efficiency of 98.87% (True good curves found / curves screened) and an acceleration of the screening process by 3.50-fold compare with the naïve approach.

The accuracy of curve classification from the embedding space based on the user-selected classes (Layer 2) was measured on an independent test set (Fig. 1D) at three different iterations. At the beginning, when only 20 curves had been labelled by the user (Fig. 2B.1), our neural network failed to correctly separate the classes contained in the train/validate subset (P1 and P2). This resulted in a unique cluster containing only some P1 curves (56/471).

After introducing more labelled traces (500, Fig. 2B.2), the model was able to correctly classify almost all P1 curves (470/471) and more than half of P2 curves (192/333). Moreover, the two selected classes were clearly separated in the embedding space and correctly clustered by a k-means model. By increasing again the number of curves visualized (1000, Fig. 2B.3) exceeding the point at which all good curves were screened, therefore including more information regarding the bad curves, the model was even able to correctly recognize Pathway 3 (i.e., the class that was not present in the training set) as good. However, due to the high degree of similarity of Pathways 1 and 3, the traces of such classes overlapped in the embedding space.

In order to correctly separate the P1 and P3 pathways in the embedding space so as to be able to recognize a class of unfolding events not present in the training set (Layer 3), we first projected the embedded data on the normal vector of the k-means separation plane. This allowed us to visualize the posterior covariance, calculated for each instance of the test set, on the ordinate. Furthermore, we separated the points using the output of the logistic regression model, differentiating between the ones classified as good (Fig. 2C.1) and the others (Fig. 2C.2). This approach provided a clear separation of all four classes (i.e. bad curves, good curves selected by the user (P1 and P2), and good curves with a previously unidentified unfolding pathway (P3)).

After showing the potential of our workflow on ideally simulated data, we tested it on real world experimental data. Experimental data poses additional difficulties such as natural signal variability within an unfolding class, and inclusion of uninterpretable complex signal patterns into the bad curves instead of just clean Gaussian noise. The bad traces spanned a wide variety of curve types representing multiple peaks and/or additional features, which made the capability of visualizing novel unselected classes more challenging, especially since some bad curves were qualitatively similar to P3. These points resulted in a slower saturation of the good curves found (Fig. 3A) in the training/validate set (196 P1 curves + 103 P2 curves = 299 good curves). In fact, after quickly sorting out the vast majority of good traces (>90%), a few more iterations were necessary to complete the screening due to the difficulty on correctly classifying the remaining ambiguous traces. Nevertheless, this was achieved after presenting 532 curves out of 1100 (48.36%), giving a 2.07-fold acceleration and a screening efficiency of 56.20%.

**Figure 3:**
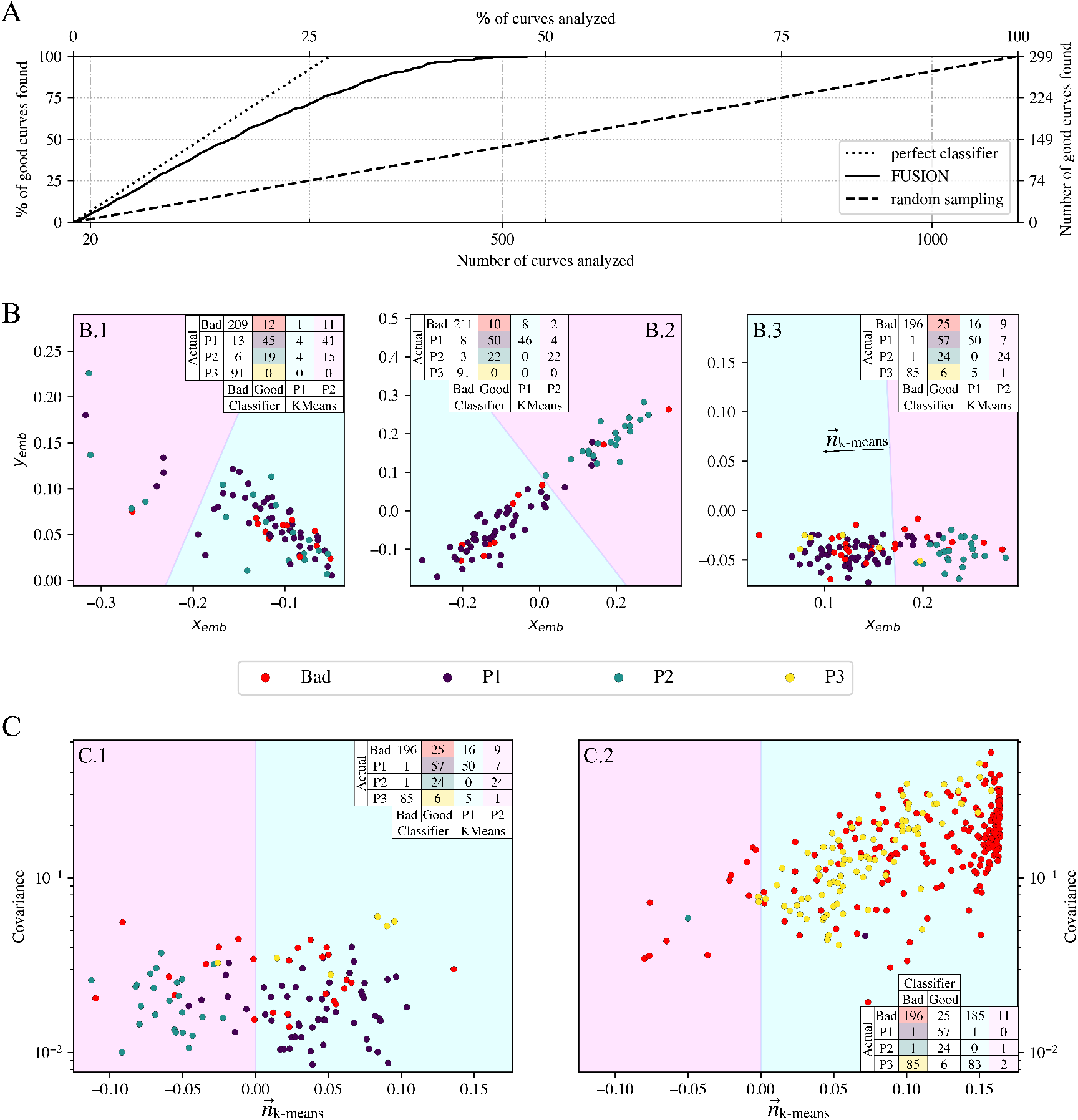
Results of the case study using experimental data. (A) Layer 1: iterative classification of good curves. The solid black line shows the result of our FUSION learning method. The dashed black line shows the theoretical random sampling. (B) Layer 2: embedding space generated at different iterations, after analyzing 20 (B.1), 500 (B.2) or 1000 (B.3) traces. The data shown come from an independent test set. Only the spectra classified as good are shown. The colors of the dots recall the true labels. The colors of the background reassemble the separation boundary of a k-means algorithm. (C) Layer 3: separation of the selected (P1 & P2), unselected pathways (P3) and bad curves using the covariance and the output of the binary classifier (layer 1) at 1000 analyzed spectra. (C.1) contains the curves classified as good, while (C.2) contains the traces classified as bad.

Regarding the embedding space, we noticed a more conservative approach of the model, especially at the beginning (20 curves visualized, Fig. 3B.1). At this stage, the model included most traces of the selected classes (P1 and P2) and no P3 curves, but also just a few bad curves enclosed in the test set. Furthermore, we observed that all classes overlaps. A necessary clarification is that, due to the probabilistic nature of running a test screening, in addition to the low number of data points at such stage, the observed behavior could not be generalized. When we analyzed the embedding at higher iterations (500 and 1000 curves screened, Fig. 3B.2 and 3B.3), we observed the absence of a large number of P3 embedded points. This suggested that the binary classifier became over-specialized in recognizing the selected good classes, ignoring the unselected ones. For this reason, we focused on the points classified negatively by the logistic regression in order to analyze the detection of P3 pathways (Fig. 3C.2). In this case, even with the posterior covariance it was not possible to completely separate P3 embedded points from the bad traces. Nonetheless, most of P3 points were located away from the main cluster of bad curves (the vertical cluster located around x=0.15 in Fig. 3C.2). Layer 3 performance could be strengthen by including additional variables to the inputs of the neural network (e.g., the scale of each curve) but, since such process would be system-specific and we focused on finding novel classes based only on the images of each trace, this would be beyond the scope of our work.

## Conclusions

In this work we presented FUSION learning, a multi-layer ML-powered iterative workflow used to accelerate the screening of AFM-SMFS data (layer 1), and to separate selected (layer 2) and unselected (layer 3) unfolding pathways only based on trace images. We validated our framework on two distinctive case studies of a mechanostable XMod-Doc/Coh biomolecular complex. The first study contained synthetic data generated with a Monte Carlo algorithm and the second concerned real experimental AFM-SMFS data. In both cases, FUSION learning accelerated the data screening process as compared to a naïve approach, was able to separate user-selected curve classes and generated promising results concerning the detection of unselected classes.

The first layer (binary classifier) could considerably help researchers in data screening, decreasing the number of traces to analyze, with an acceleration between 2- and 3.5-fold. Furthermore, our iterative process overcomes the need to blindly trust a machine learning model. The second layer (embedding space) allows separation of pathways selected by the user, giving also a view on how similar/overlapping different classes are. Layer 3 had the most challenging task to identify novel classes; however, it did not yield a completely generalizable approach. Nevertheless, in this work we presented a possible strategy (i.e., using posterior covariance) to increase the chances of finding new, undiscovered unfolding events. In general, we noticed that the applicability and the susceptibility to initial conditions of FUSION learning decreases with the increase in the layer number. The reported FUSION learning technique and analogous ML approaches can address data analysis bottlenecks and accelerate new discoveries in the field of single-molecule biophysics.

## Acknowledgement

This work was supported by the University of Basel, the Swiss Federal Institute of Technology in Zürich (ETH Zürich) and the Swiss Nanoscience Institute (SNI, project P1802). The authors declare that some text was edited using ChatGPT.^37^

## Supporting Information Available

Additional details on materials and methods, including an extended description of AFM-SMFS measurements, Monte Carlo simulations, data processing, the kernel matrix employed and the three layers of FUSION Learning algorithm. The code and the data used in this work are available free of charge on our group GitHub repository. This includes the Monte Carlo algorithm^35^ used to generate the simulated data (https://github.com/Nash-Lab/Monte-Carlo-Methods), the raw data (Zenodo DOI:10.5281/zenodo.8224236) and the script to reproduce this work (https://github.com/Nash-Lab/Fusion-Learning).

## TOC Graphic

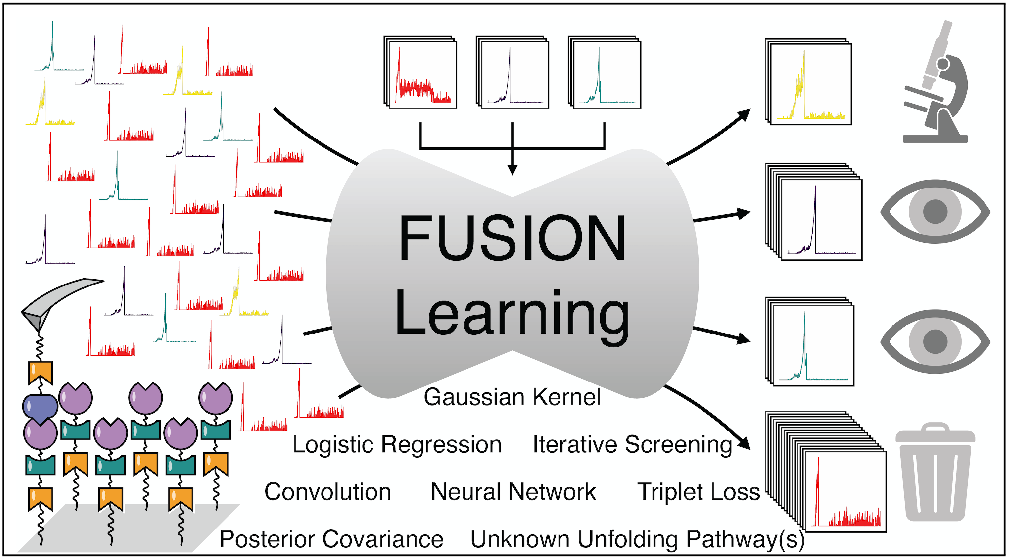

## Notes

https://github.com/Nash-Lab/

### Competing Interest Statement

The authors have declared no competing interest.

## References

(1) Müller, D. J.; Dumitru, A. C.; Lo Giudice, C.; Gaub, H. E.; Hinterdorfer, P.; Hummer, G.; De Yoreo, J. J.; Dufrêne, Y. F.; Alsteens, D. Atomic Force Microscopy-Based Force Spectroscopy and Multiparametric Imaging of Biomolecular and Cellular Systems. Chem. Rev. 2021, 121, 11701–11725.

(2) Alsteens, D.; Gaub, H. E.; Newton, R.; Pfreundschuh, M.; Gerber, C.; Müller, D. J. Atomic force microscopy-based characterization and design of biointerfaces. Nature Reviews Materials 2017, 2, 17008.

(3) Tych, K.; Žoldák, G. Stable substructures in proteins and how to find them using single-molecule force spectroscopy. Protein Supersecondary Structures: Methods and Protocols 2019, 263–282.

(4) Li, Y.; Wen, J.; Qin, M.; Cao, Y.; Ma, H.; Wang, W. Single-Molecule Mechanics of Catechol-Iron Coordination Bonds. ACS Biomater. Sci. Eng. 2017, 3, 979–989.

(5) Bernardi, R. C.; Durner, E.; Schoeler, C.; Malinowska, K. H.; Carvalho, B. G.; Bayer, E. A.; Luthey-Schulten, Z.; Gaub, H. E.; Nash, M. A. Mechanisms of Nanonewton Mechanostability in a Protein Complex Revealed by Molecular Dynamics Simulations and Single-Molecule Force Spectroscopy. J. Am. Chem. Soc. 2019, 141, 14752–14763.

(6) Dahal, N.; Nowitzke, J.; Eis, A.; Popa, I. Binding-Induced Stabilization Measured on the Same Molecular Protein Substrate Using Single-Molecule Magnetic Tweezers and Heterocovalent Attachments. J. Phys. Chem. B 2020, 124, 3283–3290.

(7) Petrosyan, R.; Narayan, A.; Woodside, M. T. Single-Molecule Force Spectroscopy of Protein Folding. Journal of Molecular Biology 2021, 433, 167207.

(8) Galvanetto, N.; Ye, Z.; Marchesi, A.; Mortal, S.; Maity, S.; Laio, A.; Torre, V.; Bassereau, P.; Aldrich, R. W.; Blanchard, A. T.; Tapia-Rojo, R. Unfolding and identification of membrane proteins in situ. eLife 2022, 11, e77427.

(9) Chetrit, E.; Sharma, S.; Maayan, U.; Pelah, M. G.; Klausner, Z.; Popa, I.; Berkovich, R. Nonexponential kinetics captured in sequential unfolding of polyproteins over a range of loads. Current Research in Structural Biology 2022, 4, 106–117.

(10) Ott, W.; Jobst, M. A.; Schoeler, C.; Gaub, H. E.; Nash, M. A. Single-molecule force spectroscopy on polyproteins and receptor-ligand complexes: The current toolbox. Journal of Structural Biology 2017, 197, 3–12.

(11) Jacobson, D. R.; Perkins, T. T. Correcting molecular transition rates measured by single-molecule force spectroscopy for limited temporal resolution. PRE 2020, 102, 022402.

(12) Ott, W.; Jobst, M. A.; Bauer, M. S.; Durner, E.; Milles, L. F.; Nash, M. A.; Gaub, H. E. Elastin-like Polypeptide Linkers for Single-Molecule Force Spectroscopy. ACS Nano 2017, 11, 6346–6354.

(13) Binnig, G.; Quate, C. F.; Gerber, C. Atomic Force Microscope. Phys. Rev. Lett. 1986, 56, 930–933.

(14) Lee, G. U.; Kidwell, D. A.; Colton, R. J. Sensing Discrete Streptavidin-Biotin Interactions with Atomic Force Microscopy. Langmuir 1994, 10, 354–357.

(15) Rief, M.; Oesterhelt, F.; Heymann, B.; Gaub, H. E. Single Molecule Force Spectroscopy on Polysaccharides by Atomic Force Microscopy. Science 1997, 275, 1295–1297.

(16) Yang, B.; Liu, Z.; Liu, H.; Nash, M. A. Next Generation Methods for Single-Molecule Force Spectroscopy on Polyproteins and Receptor-Ligand Complexes. Frontiers in Molecular Biosciences 2020, 7.

(17) Puchner, E. M.; Franzen, G.; Gautel, M.; Gaub, H. E. Comparing Proteins by Their Unfolding Pattern. Biophysical Journal 2008, 95, 426–434.

(18) Bizzarri, A. R.; Cannistraro, S. Dynamic force spectroscopy and biomolecular recognition; CRC Press, 2012.

(19) Liu, Y.; Sun, Q.; Lu, W.; Wang, H.; Sun, Y.; Wang, Z.; Lu, X.; Zeng, K. General Resolution Enhancement Method in Atomic Force Microscopy Using Deep Learning. Adv. Theory Simul. 2019, 2, 1800137.

(20) Minelli, E.; Ciasca, G.; Sassun, T. E.; Antonelli, M.; Palmieri, V.; Papi, M.; Maulucci, G.; Santoro, A.; Giangaspero, F.; Delfini, R.; Campi, G.; De Spirito, M. A fully-automated neural network analysis of AFM force-distance curves for cancer tissue diagnosis. Applied Physics Letters 2017, 111, 143701.

(21) Huang, B.; Li, Z.; Li, J. An artificial intelligence atomic force microscope enabled by machine learning. Nanoscale 2018, 10, 21320–21326.

(22) Müller, P.; Abuhattum, S.; Möllmert, S.; Ulbricht, E.; Taubenberger, A. V.; Guck, J. nanite: using machine learning to assess the quality of atomic force microscopy-enabled nano-indentation data. BMC Bioinformatics 2019, 20, 465.

(23) Daniels, A. L.; Calderon, C. P.; Randolph, T. W. Machine learning and statistical analyses for extracting and characterizing “fingerprints” of antibody aggregation at container interfaces from flow microscopy images. Biotechnology and Bioengineering 2020, n/a.

(24) Schneider, M.; Al-Shaer, A.; Forde, N. R. AutoSmarTrace: Automated chain tracing and flexibility analysis of biological filaments. Biophysical Journal 2021, 120, 2599–2608.

(25) Waite, J. R.; Tan, S. Y.; Saha, H.; Sarkar, S.; Sarkar, A. Few-shot deep learning for AFM force curve characterization of single-molecule interactions. Patterns 2023, 4, 100672.

(26) Fukushima, K. Neocognitron: A self-organizing neural network model for a mechanism of pattern recognition unaffected by shift in position. Biological Cybernetics 1980, 36, 193–202.

(27) Schroff, F.; Kalenichenko, D.; Philbin, J. FaceNet: A unified embedding for face recognition and clustering. 2015 IEEE Conference on Computer Vision and Pattern Recognition (CVPR) 2015,

(28) Huang, G.; Liu, Z.; Maaten, L. V. D.; Weinberger, K. Q. Densely Connected Convolutional Networks. 2017 IEEE Conference on Computer Vision and Pattern Recognition (CVPR). 2017; pp 2261–2269.

(29) Ben David, Y.; Dassa, B.; Borovok, I.; Lamed, R.; Koropatkin, N. M.; Martens, E. C.; White, B. A.; Bernalier-Donadille, A.; Duncan, S. H.; Flint, H. J.; Bayer, E. A.; Moraïs, S. Ruminococcal cellulosome systems from rumen to human. Environ Microbiol 2015, 17, 3407–3426.

(30) Moraïs, S.; David, Y. B.; Bensoussan, L.; Duncan, S. H.; Koropatkin, N. M.; Martens, E. C.; Flint, H. J.; Bayer, E. A. Enzymatic profiling of cellulosomal enzymes from the human gut bacterium, Ruminococcus champanellensis, reveals a fine-tuned system for cohesin-dockerin recognition. Environ Microbiol 2016, 18, 542–556.

(31) Liu, Z.; Liu, H.; Vera, A. M.; Bernardi, R. C.; Tinnefeld, P.; Nash, M. A. High force catch bond mechanism of bacterial adhesion in the human gut. Nature Communications 2020, 11, 4321.

(32) Liu, H.; Liu, Z.; Sá Santos, M.; Nash, M. A. Direct Comparison of Lysine versus Site-Specific Protein Surface Immobilization in Single-Molecule Mechanical Assays**. Angew. Chem. Int. Ed. 2023, n/a, e202304136.

(33) Deng, J.; Dong, W.; Socher, R.; Li, L. J.; Li, K.; Fei-Fei, L. ImageNet: A large-scale hierarchical image database. 2009 IEEE Conference on Computer Vision and Pattern Recognition. 2009; pp 248–255.

(34) Rasmussen, C. E.; Williams, C. K. I. In Gaussian Processes for Machine Learning ; Dietterich, T., Ed.; The MIT Press, 2006.

(35) Liu, H.; Liu, Z.; Yang, B.; Lopez Morales, J.; Nash, M. A. Optimal Sacrificial Domains in Mechanical Polyproteins: S. epidermidis Adhesins Are Tuned for Work Dissipation. JACS Au 2022, 2, 1417–1427.

(36) https://github.com/NashLab/Monte-Carlo.

(37) OpenAI, GPT-4 Technical Report. 2023.

