## Supplementary Information for "Iterative Machine Learning for Classification and Discovery of Single-molecule Unfolding Trajectories from Force Spectroscopy Data"

### Methodology

#### AFM SMFS measurement

Experimentally, AFM-SMFS was conducted on a Force Robot AFM (JPK instruments, Berlin, Germany) as described before.[1] In brief, the Coh and XMod-Doc molecules were site-specifically and covalently conjugated onto the Biolever mini AFM cantilevers (Bruker, Billerica, MA, USA) and a glass surface respectively, using a Sfp-catalyzed surface conjugation method. Cantilever spring constants (ranging from 0.07 to 0.1 $Nm^{-1}$) were calibrated using the contact-free method. A forced pulling process was applied under a constant pulling speed of 400 nm/s. The force-extension traces were recorded from which the rupture of the complex could be analyzed. The rupture forces and corresponding force loading rate were fit with a theoretical model (Bell-Evans[2,3]) to extract energy landscape parameters. Based on the energy profile describing the stochastic dissociation of the complex, simulated force-extension traces were generated using a Monte Carlo simulation based on Kramers' theory.

#### Monte Carlo simulation

Monte Carlo simulation[4,5] was conducted to generate the simulated SMFS traces based on the energy profiles of the XMod-Doc/Coh complex. To start with, the XMod-Doc/Coh complex was randomly assigned a binding mode to be either high force binding mode (80%, Pathways 1 and 2) or low force binding mode B (20%, Pathway 3). Then a series of force values F(ti) was generated on an evenly distributed extension axis x(ti) using a worm-like chain (WLC) model.[6] A constant speed protocol was applied on the tandem molecular system including the XMod-Doc/Coh complex and the AFM cantilever with a spring constant of 91 pN/nm. During each time step $\Delta$t, the probability of XMod-Doc/Coh rupture and XMod unfolding was calculated using the force dependent off-rate obtained from a Bell-Evans model.[2,3] The event probabilities (P(F)) were compared to a random number between zero and unity. If the random number was smaller than P(F), the complex rupture (or XMod unfolding) event was considered to have occurred within that time step, and the corresponding force was recorded as the rupture.

#### Data Processing

We used two AFM-SMFS datasets, the first generated from Monte Carlo simulations[4,5] and the second containing AFM experimental data.[1] Each of the raw curves (Fig. 1B) was down-sampled to 600 points, the sign of the force variable was inverted ($f = -f$), clipped in the positive domain ($0 \leq f < \infty$) and normalized to its maximum ($f = f/fmax$). The resulting 200x200 black and white images (Fig. 1C) were pre-processed to be fed to a pre-trained CNN (Densenet121[7]). This included: i) resizing every image to 243x243x3 using the nearest-neighbor interpolation algorithm; ii) pre-processing the images using the appropriate input function. Both operations were carried using the built-in functions of the TensorFlow library.[8] The TensorFlow implementation of Densenet121, pre-trained on ImageNet dataset[9] was utilized as a feature extractor. The fully connected layer on top of the network was excluded, resulting in a vectorial output containing 1024 values (features) per each input images.

#### Kernel Matrix

The outputs of Densenet121 (x) was utilized to calculate an Euclidian distance matrix (eq. 1) and a gaussian kernel similarity matrix (eq. 2).

$$d_{i,j} = d(x_i, x_j) = |x_i - x_j|_2^2 \tag{1}$$

$$K_{i,j} = K(x_i, x_j) = exp\left(-\frac{d_{i,j}}{2\sigma^2}\right) \tag{2}$$

Where $2\sigma^2$ is the kernel scale hyperparameter. The rows of such matrix correspond to the data points used to evaluate the model ($x_i$, training/validation/test), while the columns resemble the training data ($x_j$).

#### First Layer

A logistic regressor from sci-kit learn library was built on top of the gaussian kernel matrix for the first layer of our model. FUSION algorithm (Fig. 1E) was initialized by randomly selecting 4 images from the train/validate set. During each following iteration, 4 new traces were added to the train/validate subset accordingly with the highest output of the logistic regression tested on the remaining curves (unscreened data, Fig. 1E, main text). At each iteration, the train/validate subset was split in a training (80%) and validation (20%) subsubset. After that, the traces were up-sampled to match the highest binary class present in the corresponding subsubsets (good/bad). The kernel scale was optimized via grid search using a unitary spaced log-2 scale (from $2^{-2}$ to $2^{13}$).

#### Second Layer

A fully connected neural network from TensorFlow library was built on top of the gaussian kernel matrix to generate an embedding space in which to visualize the data (Layer 2). This neural network consisted of a single output layer with 2 neurons and no hidden layers and it was trained with TripletHardLoss[10] function from the TensorFlow Addons repository. The network was initialized after each iteration and trained for a maximum of 10'000 epochs with an EarlyStopping (patience=50) and a ReduceLROnPlateu (factor=0.1, patience=10, cooldown=5) callbacks. Note that the training was performed excluding the curves selected as Bad. Similarly to what was done for the First Layer, the train/validate subset was again divided (80%/20%) and the curves were up-sampled accordingly to the highest pathway present (P1/P2). The kernel scale was fixed to the value found during the optimization of the First Layer. To better characterize the embedding space, we utilized a k-means clustering function from sci-kit learn library using two clusters (P1/P2). The Second Layer was always tested on an independent set (Fig. 1D, main text), which contained also P3 curves. The datapoints visualised (Fig. 2B.1-3 and Fig. 3B1-3, main text) were the ones which passed the binary classifier filter (the logistic regression of layer 1).

#### Third Layer

In order to further separate the unselected pathway (P3) from the ones known by the user (P1 and P2), we started by separating the test data accordingly with the output of the logistic regression of the First Layer. Then, we calculated the posterior covariance[11] (eq. 3) for each data point contained in the test set.

$$cov = diag\left(K(X_*, X_*) - K(X, X_*)^T \left(K(X, X) + 10^{-8} I\right) K(X, X_*)\right) \tag{3}$$

Where $K$ is the gaussian kernel (eq. 2), $X$ is the matrix containing all data in the train/validate set and $X_*$ is the matrix containing all test data. Finally, we plotted the posterior covariance against the embedded test data projected on the normal vector of the k-means separation line.

### References


(1) Liu, Z.; Liu, H.; Vera, A. M.; Bernardi, R. C.; Tinnefeld, P.; Nash, M. A. High force catch bond mechanism of bacterial adhesion in the human gut. *Nature Communications* **2020**, *11*, 4321.

(2) Evans, E.; Ritchie, K. Dynamic strength of molecular adhesion bonds. *Biophysical Journal* **1997**, *72*, 1541–1555.

(3) Bell, G. I. Models for the Specific Adhesion of Cells to Cells. *Science* **1978**, *200*, 618–627.

(4) Liu, H.; Liu, Z.; Yang, B.; Lopez Morales, J.; Nash, M. A. Optimal Sacrificial Domains

in Mechanical Polyproteins: S. epidermidis Adhesins Are Tuned for Work Dissipation. *JACS Au* **2022**, *2*, 1417–1427.

(5) https://github.com/NashLab/Monte-Carlo.

(6) Bustamante, C.; Marko, J. F.; Siggia, E. D.; Smith, S. Entropic Elasticity of lambda-Phage DNA. *Science* **1994**, *265*, 1599–1600.

(7) Huang, G.; Liu, Z.; Maaten, L. V. D.; Weinberger, K. Q. Densely Connected Convolutional Networks. 2017 IEEE Conference on Computer Vision and Pattern Recognition (CVPR). 2017; pp 2261–2269.

(8) Abadi, M. et al. TensorFlow: Large-Scale Machine Learning on Heterogeneous Systems. 2015; `https://www.tensorflow.org/`, Software available from tensorflow.org.

(9) Deng, J.; Dong, W.; Socher, R.; Li, L. J.; Li, K.; Fei-Fei, L. ImageNet: A large-scale hierarchical image database. 2009 IEEE Conference on Computer Vision and Pattern Recognition. 2009; pp 248–255.

(10) Hermans, A.; Beyer, L.; Leibe, B. In Defense of the Triplet Loss for Person Re-Identification. 2017.

(11) Rasmussen, C. E.; Williams, C. K. I. In *Gaussian Processes for Machine Learning*; Dietterich, T., Ed.; The MIT Press, 2006.
